# VIRALpre: Genomic Foundation Model Embedding Fused with K-mer Feature for Virus Identification

**DOI:** 10.1101/2024.11.12.623150

**Authors:** Zanyi Wang, Qinze Yu, Yu Li

## Abstract

Virus, a submicroscopic infectious agent, influences all life forms. Identifying viral sequences is essential to understand their biological functions and then analyze their impacts on public health, and the development of microbial communities. For its significance, tools are developed based on various mathematical methods and algorithms. However, previous methods struggle to identify viral sequences, especially short contigs accurately since the limited information and small-scale close-set dataset. Here we propose VIRALpre, a hybrid framework combined with genomic foundation model (GFM) embedding and K-mer feature of sequences to precisely recognize viral genomic fragments. VIRALpre is empowered by the generalization competencies of GFMs, which have proven their strength in various downstream tasks, thanks to newly established large-scale training databases and Attention mechanism. On the other hand, K-mer features provide additional biological information to bridge the limitation of GFMs in classification tasks. Comprehensive experimental results demonstrate that VIRALpre significantly outperforms all the previous methods on virus identification performance by 4% in accuracy. To prove that this model is qualified when facing unique contigs to training data, BLASTn-based similarity cut-off test(setting e-value as 10^−5^) is done and it achieves about 10% F1-score improvement. More than well-built test datasets, new zero-shot cross-dataset tests on benchmark datasets sampling from natural environments are conducted, VIRALpre performs identify almost most viral sequences while keeping a very low False Positive Rate. Based on these solid experiments, VIRALpre has the ability to manage short-contig virus identification by truly learning the distinctions of viral sequences and hopefully act as an adviser to promote virus-related research.

## 1 Introduction

The virus is widespread across almost all ecosystems and can affect all life forms. Being an important agent in nature, viruses can influence all individuals and then impact the evolution of the biological community. Identifying virus sequences accurately has biological and medical significance since it helps to track the spread of diseases, understand the ecological dynamics, enable the development of antiviral drugs, and implement effective public health measures to contain outbreaks.

Many tools and methods have been used to identify specific species sequences. Previous methods can be divided into three groups [1], homology-only, machine learning, and deep learning methods. Homology tools compare the similarity between the candidate sequence and sequences in the database. For example, MetaPhinder [2] uses BLASTn [3] to calculate the e-value [4] as the criteria for classifying sequences. These methods achieve good performance, however, they rely on the database and struggle when they face sequences unsimilar to reference sequences. Machine learning methods, utilize machine learning algorithms, such as HMM [5](hidden Markov model) and Random Forest [6], to classify metagenomic data. Conversely, these methods rely on long sequences as input to model the relationship between nucleotide or protein markers as indicators to instruct classification. Take Virsorter2 [7] as an example, its performance increases when sequence lengths are longer than 3k byte pairs. Other methods, such as VIBRANT [8] and Genomad [9], use protein markers to identify viral sequences. These methods need the sequences to have enough proteins and coding regions, which restrict their performance, especially in short viral sequence analysis. Deep learning tools apply neural networks as their backbones. DeepVirFinder [10] and PPR-Meta [11] use CNN [12] (convolutional neural network) to identify viral sequences and Seeker [13] employs LSTM [14] to handle genomic data. CNN is good at capturing global and regional dependence and LSTM can learn long and short-term relationships between byte pairs, however, these deep learning methods lack generalization performance because of the limitation of close-set training databases. Specifically, these models perform decreasingly when managing unique sequences to the training dataset and gain low precision if negative samples are from species different from the training dataset.

Recently, large Language Models (LLMs), such as Gpt-4 [15], have achieved extraordinary performance in almost all downstream in natural language process based on Transformer architecture [16]. In genomic research, more and more genomic sequences are available to train deep-learning models. In this stage, Genomic Foundation Models (GFMs) [17, 18], resorting to their generalization performance and large-scale pretrained data, can achieve stunning results in many downstream tasks, including contig classification. Previous methods, such as ViraLM [19], take full advantage of DNABERT-2 [20]. However, fine-tuning foundation models fully relies on the embedding result of LLM, and neglects the representation of original contigs. Meanwhile, different from natural language, biological information is more intricate and plentiful. Genomic foundation models are still there from completely understanding the representation of metagenomic data and making accurate classification.

For these problems, we propose VIRALpre, a K-mer [21]feature fused foundation model based classifier. The K-mer feature module in VIRALpre serves as a client to compress input sequences into a fixed-length vector that contains sufficient virus-particularity information to promote the result in identifying viral contigs. Besides the two modules, a fusion module is designed to combine the output of these two parts and get reliable results. A feature fusion module is proposed based on MLP [22] (multi-layer perceptron) and CNN (convolutional neural work). The two feature vectors go through MLP first to normalize. After that, convolutional layers and pooling layers [23] function to gain the dependency of these two normalized information. In the end, the result is projected by a linear layer to distinguish whether the input contig is a vial sequence or not.

In five-fold cross-validation, VIRALpre is the best model and achieves almost 4% increase in accuracy. To test how these models perform when encountering unsimilar input sequences to training data, BLASTn is used to filter similar conditions in the test dataset. For this similarity cut-off dataset, VIRALpre gets the best performance which enhances accuracy by about 10% compared with other models. More than the RefSeq database, third-party datasets are used to get comprehensive test results. Three benchmark datasets sample data from natural environments are used to test generalization ability and verify if VIRALpre can be qualified to discover new viral sequences. In this test, VIRALpre identifies a great number of positive sequences and keeps a very low False Positive Rate.

The code is available at https://github.com/xmz111/VIRALpre.

## 2 Methods

### 2.1 Overview of Method

As depicted in Figure 1 and detailed below, the proposed framework comprises three major modules: genomic foundation model, Kmer feature, and feature fusion module.

**Figure 1:**
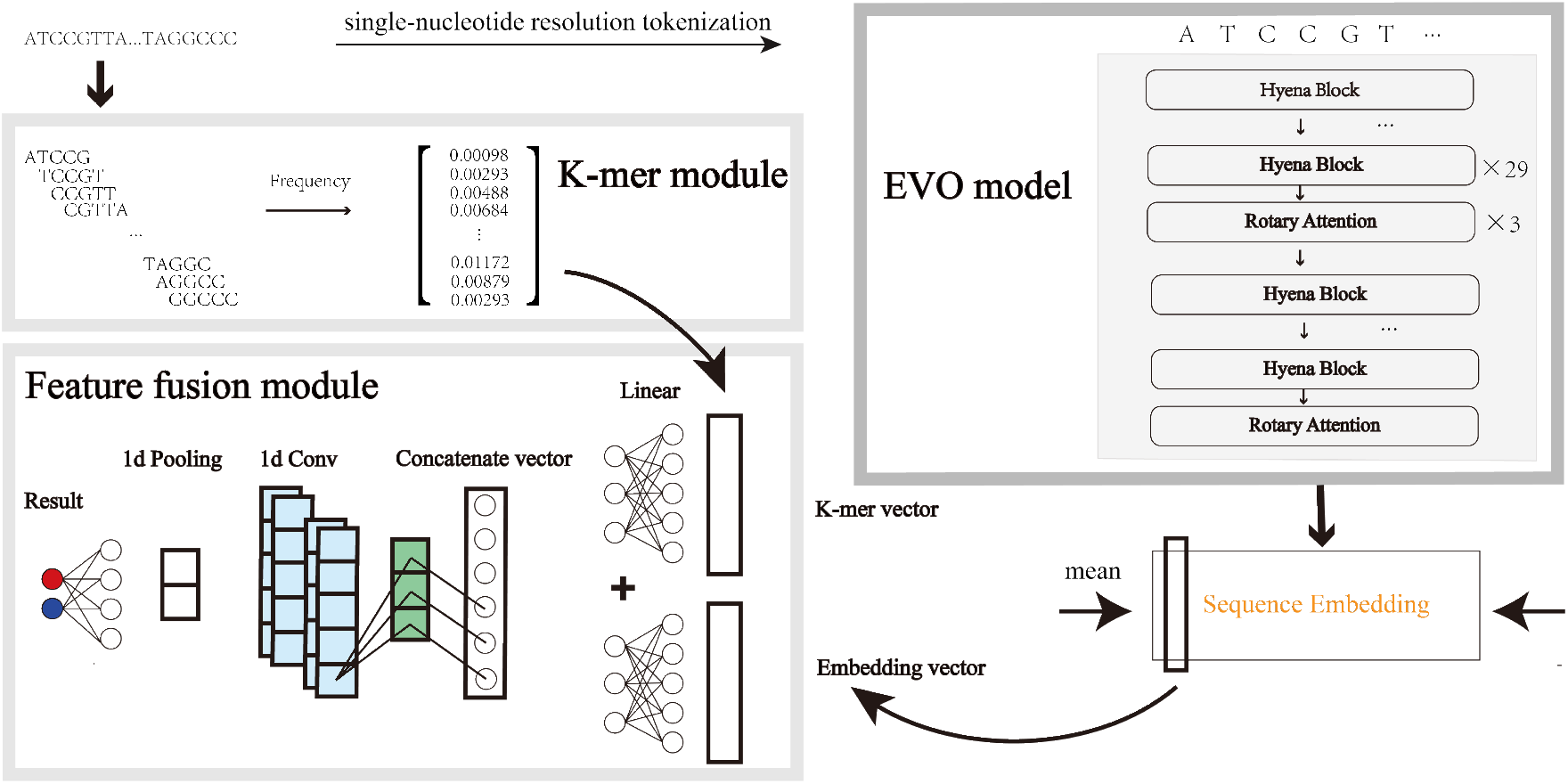
Overview of the pipeline of VIRALpre: our model takes advantage of genomic foundation models and K-mer features of input sequences. The input goes through the EVO [24] model and embedding is extracted and compressed. Meanwhile, K-mer features are calculated. In the end, the embedding and K-mer vectors are fused by MLP and CNN to obtain the final results.

Given a sequence, it will be input into the foundation model module and K-mer module seprately. For the foundation model part, the sequence goes through single-nucleotide resolution tokenization and input into EVO model. The embedding of the sequence is extracted and compressed into the embedding vector. At the same time, the appearance frequencies of various K-mer components are gained. Eventually, these two vectors are fused by the feature fusion module.

### 2.2 Genomic foundation model module

Genomic foundation models are pre-trained by a large number of genomic data and make great achievements on many downstream tasks. Recently, EVO [24] is proposed for its novel architecture, StripedHyena [25] and large-scale pre-trained dataset-OpenGenome, so it is selected as the used genomic foundation model.

The whole model is comprised of 29 hyena blocks and 3 attention blocks. For Hyena block, short convolutions, long convolutions and gating models long sequences. Suggests that the input tensor is *x*, convoultional kernel is *h*, convolutional projection can be expressed as:

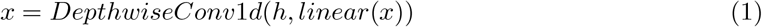

Then *x* is split into *q, k, v* and input into the attention block, the attention mechanism can be expressed:

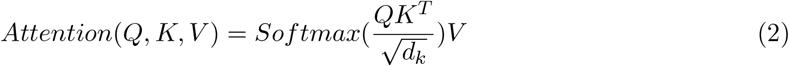

In this equation, *d*_*k*_ is the dimension of *K, Q, K, V* are obtained as example:

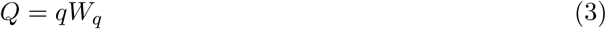

Among the equation, *W*_*q*_ is a trainable matrix.

Because the EVO model doesn’t have a CLS token, the embedding result is used to make the classification. Suppose that the embedding result can be expressed as: *E*^*d×L*^, *d* is the dimension of the EVO model, which is 4096, *L* is the input length of sequences. To replace the CLS token, the mean value of the embedding matrix in length dimension is calculated as the foundation model feature vector, whose length is *d*.

### 2.3 K-mer feature module

The K-mer feature is a statistical feature that can reflect the nucleotide component of a sequence and the short-term dependency of nucleotide. For each sequence, K-mer feature compresses the biological information into a vector no matter how long it is. Virus identification tools which based on deep-learning method model sequences by tokenization them into fixed-length tensors. This method utilizes the computational competence of GPUs, however, the original semantic information of genomic sequences will lose and change because of padding and truncating. By introducing the K-mer feature, the model receive the whole biological information of sequences which further improves the model performance in the metagenomic dataset.

Suppose that a genomic sequence is a string whose length is *L*. To get k-mer features, sequences are broken down into smaller segments (K-mers) of fixed length K. Then, analyze the frequency and distribution of these k-mers in the sequence. In VIRALpre, K is set as 5, and the K-mer vector has dimension 4^5^ = 1024.

### 2.4 Feature fusion module

To effectively fuse these features, CNN based feature fusion module is proposed. Compared with MLP alone, CNN based module better captures the dependency between these features and easily manages input variance issues to achieve better performance in comprehensive tests. Compared with using LSTM, CNN is faster and utilizes GPU computational ability. Moreover, LSTM struggles to manage heterogeneous features that are combined artificially and are less stable due to overfitting issue.

After the input goes through two modules, an embedding and K-mer vector, namely 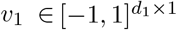 and 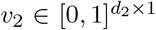 are got, which contain two different biological information and can assist to make decision. For EVO, *d*_1_ is 4096 and for 5-mer feature, *d*_2_ is 1024. In this module, two MLPs are used to project the two dimensions into the same first to promote fusion, and then concatenate in dimension 0:

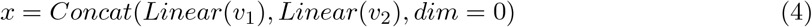

After this function, the two features are normalized, which facilitate the convolutional layers to fuse the two heterogeneous vectors. The output of the linear function is a vector *x* ∈ [−1, 1]^*d×*1^, where *d* is set as 128, and the 0th dimension of the concatenated vector is 2 × *d*. To capture the global and regional connection of this vector, convolution and maxpool layers are used:

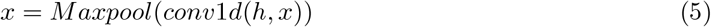

In this function, *h* is the convolutional kernel and the size is set as 3. In the first convolutional layer, 64 kernels are used to extract the inner features in the concatenated vector. Pooling is used to decrease the dimension and alleviate overfitting issues. The next convolutional layer is same as the this layer, but the number of kernels is reduced to 32. After two convolutional layers, a Linear function is used in the last layer to calculate the probability that the input sequence is a virus.

## 3 Results

### 3.1 Dataset and data process

Training data are all from the RefSeq dataset. We include all viral sequences collected by August, 2024 as positive samples and bacteria, fungi, plasmid, and archaea comprise negative datasets. Moreover, positive and negative samples are balanced, so negative datasets retain nearly the same number as positive sequences by random selection. The input sequences are filtered to keep those sequences that contain nucleotides other than A, T, C, and G. As mentioned above, our model is built to mainly fill up the gap of previous tools in short-contig identification, so all original sequences are split into short contigs as input. Specifically, their lengths are from 900 bps to 1100 bps. After that, the test dataset is determined by randomly dividing almost 20% of the whole dataset. Training data are used to build a BLASTn database, which represents information the model has learned. Then, similarity cutoff proceeds for the test data to eliminate similar sequences in the test dataset, and e-value is set as 1 × 10^−5^. After that, almost half of sequences are reserved to make up the similarity cut-off test dataset.

In test using third-party dataset, long sequences are split into short contigs whose lengths are 1024 first. For each contig, viral scores are calculated. Finally, the average of these scores for one sequence will be the result that a sequence is a virus or not.

For cross dataset test, two simulated RefSeq datasets [26] are selected to test the performance of models when they encounter unsimilar mock short contigs that have the mean length 2000, but are homologous. Moreover, three benchmark real-world datasets [1], which are split into 8 sub-dataset, are used to test the generalization competence of our model, including samples from seawater, soil and gut. Samples from these environments are filtered through 0.22 µm filters to obtain microbial- (*>*0.22 µm) and viral-enriched fractions (*<*0.22 µm). Then, DNA was separately extracted, purified, quality control and combined into longer sequences. Detailed information is available in Table 1.

**Table 1:** Details about datasets.

| Dataset | Source | Split | Length region | Positive amount | Negative amount |
| --- | --- | --- | --- | --- | --- |
| Train | RefSeq [26] | True | 900-1100 bps | 400878 | 427254 |
| Test | RefSeq | True | 900-1100 bps | 66780 | 75134 |
| Cut-off test | RefSeq | True | 900-1100 bps | 18063 | 25432 |
| Simulated | Third-party | False | 1500-2500 bps | 61550 | 61550 |
| Seawater | Third-party | False | 1500-5 * 10<sup>5</sup> bps | 2.3 * 10<sup>5</sup> | 4.9 * 10<sup>5</sup> |
| Soil | Third-party | False | 1500-1.7 * 10<sup>5</sup> bps | 1.7 * 10<sup>5</sup> | 1.1 * 10<sup>5</sup> |
| Gut | Third-party | False | 1500-1.6 * 10<sup>5</sup> bps | 8 * 10<sup>3</sup> | 3.5 * 10<sup>4</sup> |

### 3.2 Test result

In the test, other deep-learning-based virus identification tools (DeepVirFinder, ViraLM, Seeker) are re-trained using the same data as VIRALpre to keep fairness. The backbones of these three tools are CNN, DNABERT-2, and LSTM. For these four models, five-fold validation based on the training dataset is deployed first. Then, these models are trained using the whole training dataset and test on the test dataset. To prove whether models can identify unsimilar sequences to the training dataset, BLASTn is used to filter similar contigs in the test dataset. The positive and negative sequences are used to build the BLASTn database. Then, the e-value [4] is set 10^−5^ as a threshold to sift the sequences which are similar to the training dataset. The sifted dataset is used to test these models next.

In third-party dataset, these four same-daatset-trained deep-learning-based models, trained machine-learning based tool VirSorter2 and database-based method MetaPhinder are added. By the way, MetaPhinder uses training dataset as its BLASTn database to make classification.

Setting viral sequences as positive samples and other contigs as negative data. True positive rate (TPR, also known as sensitivity and recall), false positive rate (FPR), precision, accuracy, and F1 score are metrics to evaluate the performance of models. The equations are below.

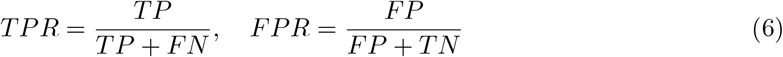

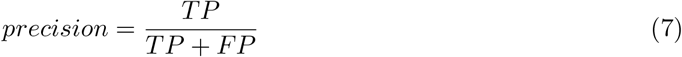

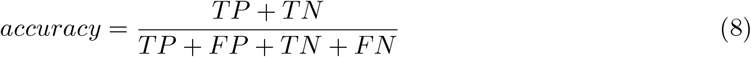

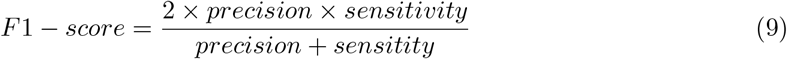

Based on the RefSeq dataset, Figure 2 shows results including Five-fold cross-validation, test consequences, results of similarity cut-off test dataset, and test on RefSeq simulated dataset. In five-fold cross-validation (a), VIRALpre achieves the best result. After training these models using the whole training dataset, the test accuracy of VIRALpre is 94%, compared with 91% accuracy achieved by DeepVirFinder, the second best model. When test on the similarity cut-off dataset, in Figure 2 (b), VIRALpre outperforms other models a lot, which is attributed to the generalization ability of foundation model and abundant input features. Although performances of all tools decrease, genomic foundation model-based model, the accuracy of VIRALpre and ViraLM reduce about 2%, compared with 10% decrease in DeepVirFinder and Seeker, which reveals the limitation of deep-learning based models in generalization competence.

**Figure 2:**
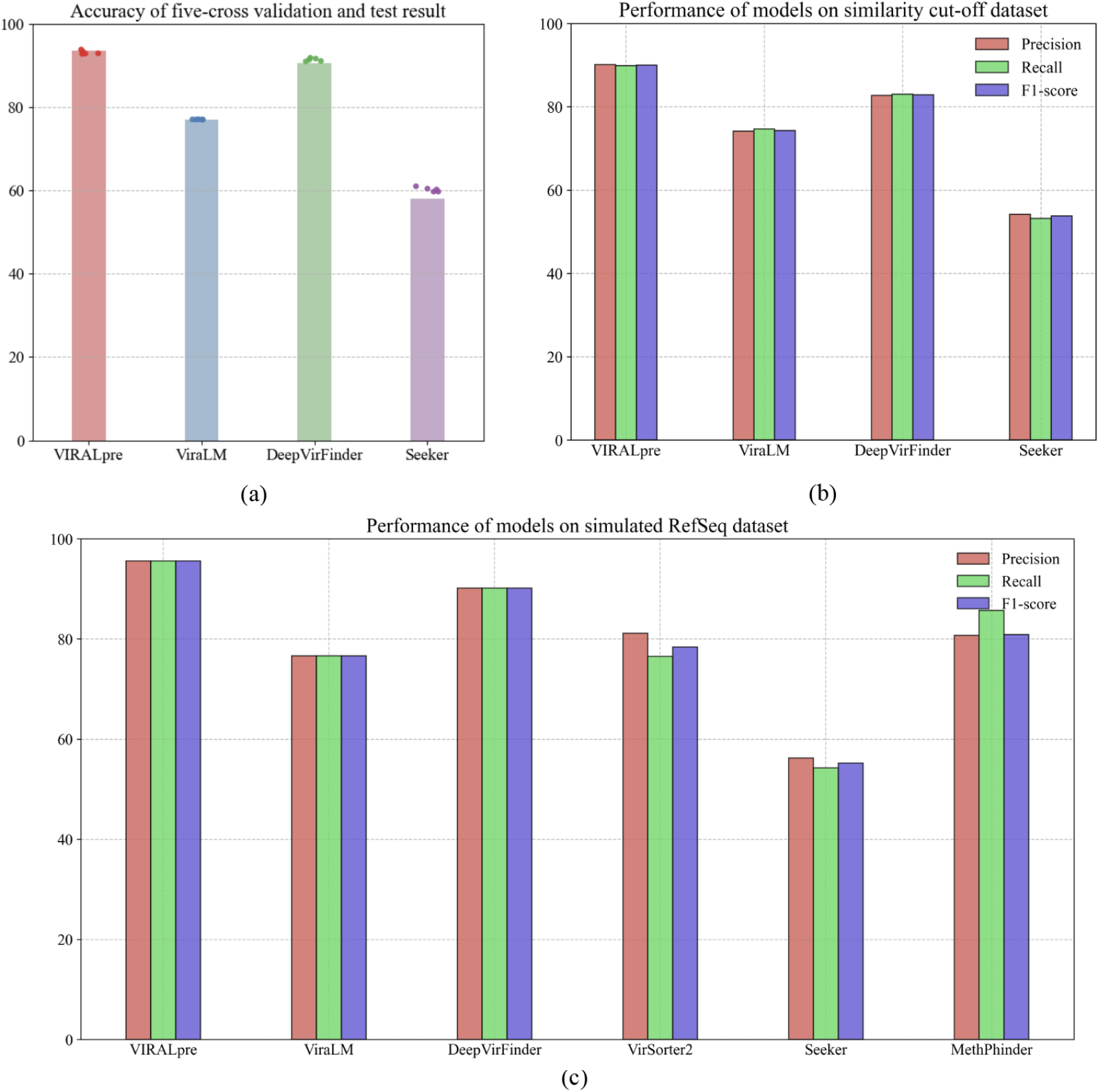
VIRALpre achieves the SOTA results in the RefSeq-related datasets. **a.** Test and five-fold cross-validation results on the RefSeq dataset, accuracy is the metric. **b**. BLASTn is used to cut off sequences in the test datasets similar to the training dataset. Precision, recall, and F1-score are metrics to do similarity cut-off test. **c**. Test various virus identification tools on third-party simulated RefSeq dataset.

Different from datasets directly from RefSeq, third-party RefSeq-simulated datasets are used to test models. More than four trained models, machine-learning based VirSorter2, which is trained on RefSeq database, and database-based tool MetaPhinder are used to prove the great performance of VIRALpre. The whole dataset contains virus-simulated sequences as positive samples and bacteria-simulated sequences as negative data. The dataset is nearly balanced. In this test, VIRALpre still achieves the highest scores in precision, recall and F1-score. The scores of these three metrics are almost the same, which indicates the stability of VIRALpre’s performance. For VirSorter2, the recall value is low, which means it identify fewer viral sequences. MethPhinder, whose database is built using a training dataset, gets great performance in identifying viral contigs since sequenes in the database is similar to positive samples, however, makes a lot of mistakes when identifying bacteria contigs.

Real-world datasets are established using samples in environments. The lengths of sequences are various. The results are in Figure 3. In the seawater dataset, VIRALpre is the best tool which identify the most viral sequences, while making a few mistakes. For the soil dataset, VIRALpre’s TPR is on third place, lower than DeepVirFinder and VirSorter2. However, its FPR is the lowest. What is interesting is that the FPR of MethaPhinder is higher than TPR, which indicates that negative samples in this dataset are more similar to positive data in the training dataset according to BLASTn. When it comes to the gut dataset, the TPR of all models decreases under 40%. In this test, VIRALpre doesn’t discover a lot of viral sequences compared with other tools. Nonetheless, the FPR is absolutely the lowest, proving the identification stability of VIRALpre.

**Figure 3:**
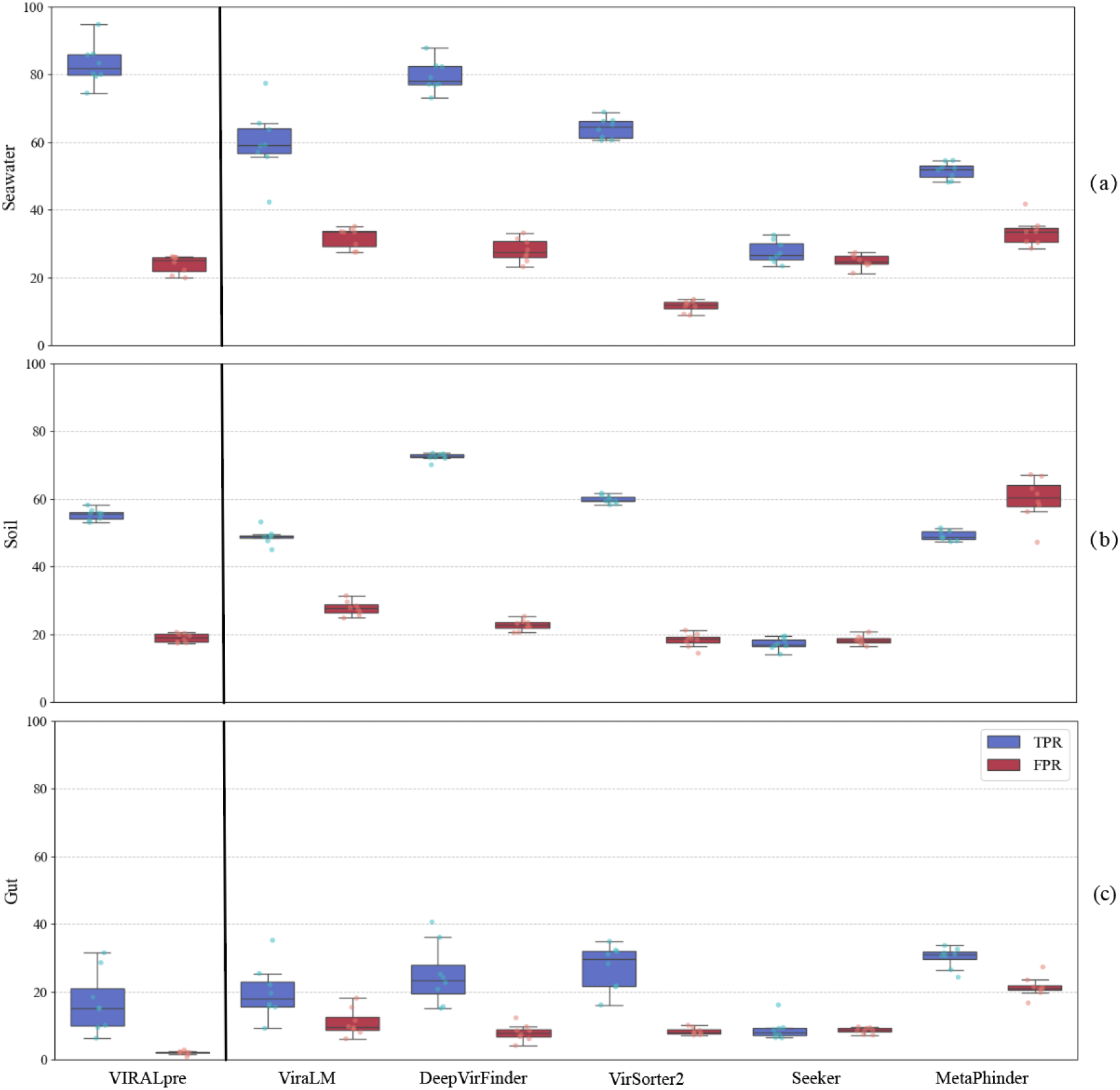
VIRALpre identifies a large number of viral sequences and remains low FPR in the real-world dataset. TPR and FPR are used to indicate the performance that models identify strange contigs. **a.** Test on samples from seawater. **b**. Test on soil sequences. **c**. Test on contigs from the gut.

### 3.3 Ablation experiment

To prove the superiority of the VIRALpre’s pipeline, the 2-D projections of the output of GFM. Figure 4 (a) and K-mer module (b) are obtained based on the T-sne [27] algorithm. The input data are from four species, virus, plasmid, bacteria, and archaea. There are 20000 contigs of each species input to each module. To compare the clustering results of VIRALpre modules, the input sequences are encoded into one-hot vectors (c).

**Figure 4:**
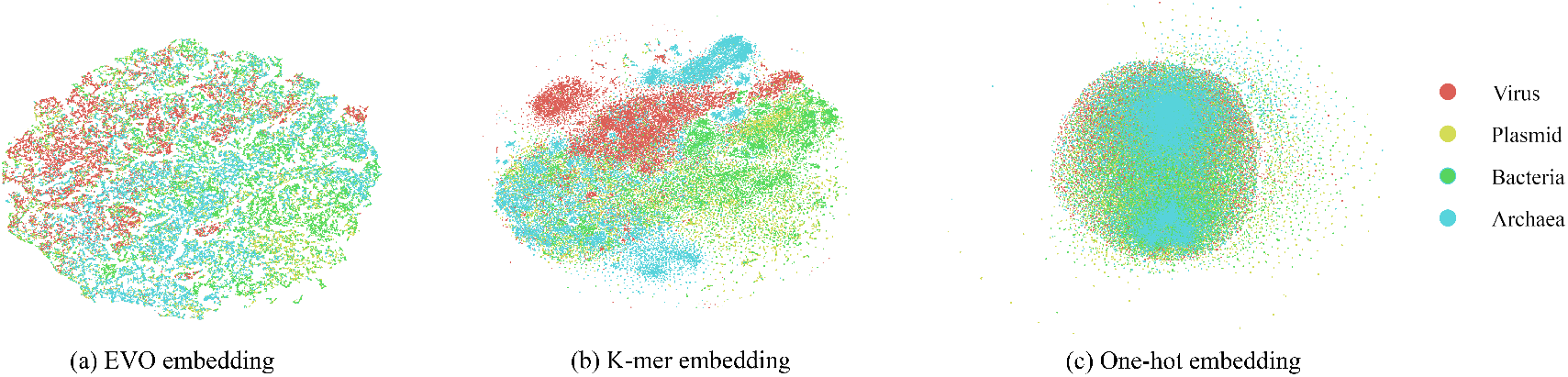
GFM and K-mer feature distinguish sequences from different species, but not enough to clearly identify viral sequences. Clustering effects of feature extraction modules managing different species sequences based on T-sne algorithm (inputs are processed vectors) are shown: **a.** EVO embedding compressed vectors. **b**. K-mer vectors. **c**. One-hot encoding vectors.

One-hot embedding doesn’t have an obvious cluster boundary and too many data points are in the central area. For EVO embedding and K-mer embedding, it is clear that the data points are separated, especially viral sequences, however, for these two modules, the clustering results are not ideal enough. To further analyze the concrete difference each module makes, VIRALpre is split and combined into 4 sub-models, foundation model only, K-mer only, GFM embedding, and K-mer features fused by simple MLP, and VIRALpre, which replaces MLP with the feature fusion module. Training datasets are inputted into these models and use test datasets to test the results.

In Figure 5 (a), only the K-mer feature or GFM achieves relatively poor results in viral identification. When combining these modules together using MLP, the performance increases by nearly 10%, which indicates the effectiveness of fusing these modules. When replacing MLP to the feature fusion module, the performance increases by about 8%, manifesting that this module is important in fusing heterogeneous features.

**Figure 5:**
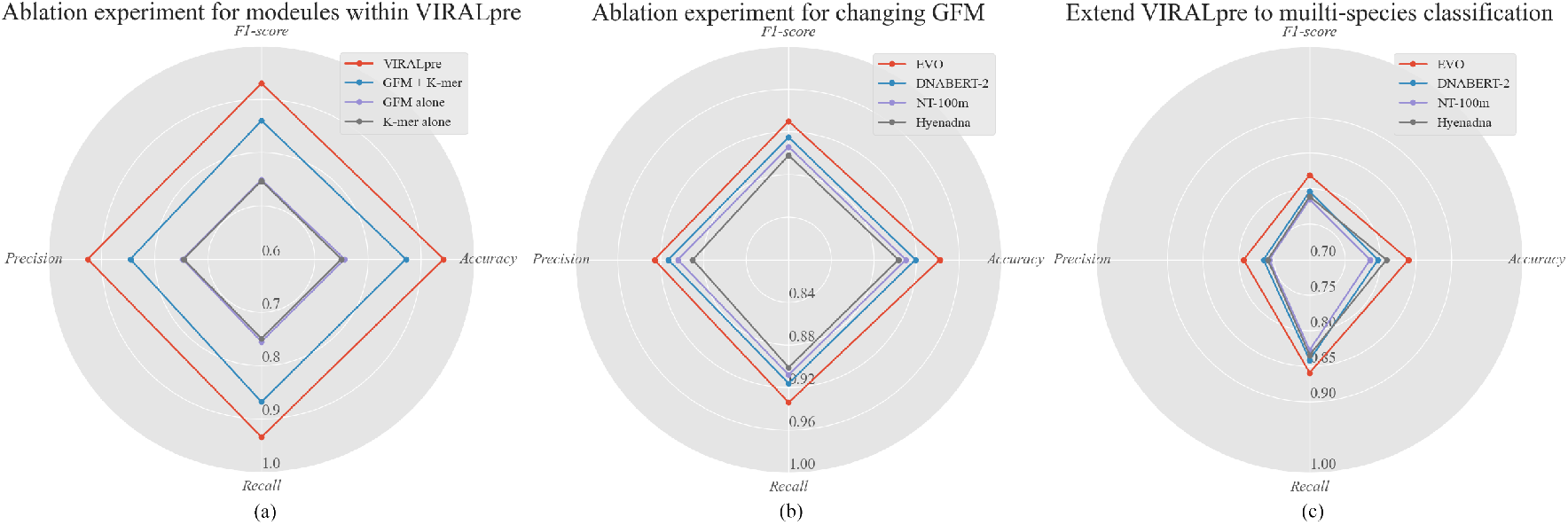
Ablation results and the improvement of VIRALpre. F1-score, precision, recall, and accuracy are chosen as the metrics. **a.** Test module combinations of VIRALpre. **b**. Binary classification results from VIRALpre training framework where foundation model is changed. **c**. EXtend the framework of VIRALpre to multi-species classification task, of replaced foundation model VIRALpre.

To analyze the function of GFMs, further than utilizing EVO [24] as the backbone, another GFMs are used as well. The experiments use four pre-trained genomic foundation models, namely Hyenadna [28], DNABERT-2 [20], Nucleotide Transformer [29], and EVO as the backbone. Detailed information is available on Table 2.

**Table 2:** Details about different foundation models.

| Models | Parameters | Pre-trained data | Architecture | Embedding dim |
| --- | --- | --- | --- | --- |
| Hyena-medium | 6.6M | human-reference genome [30] | Hyena [31] | 256 |
| DNABERT-2 | 117M | multi-species [32] | Bert [33] | 768 |
| NT-100m | 100M | multi-species | Transformer [16] | 512 |
| EVO | 7B | Open-Genome [24] | Striped Hyena [25] | 4096 |

Training data remains the same as the training stage. To extend viral identification further, a multi-classes classification model is trained by splitting negative samples into four original species, namely plasmid, bacteria, fungi, and archaea. To alleviate the unbalanced dataset issues, the weights of these classes are added into cross-entropy loss to keep fairness.

The result is shown in Figure 5 (b), (c). The selection of EVO attains the best results, which has almost 3% improvement compared with the second best backbone-DNABERT-2. The boost of performance attributes to larger pre-trained data and higher embedding dimension of EVO. In multi-classes classification, which is harder than binary classification, the recall values are higher than precision a lot because of the unbalanced dataset. In this test, VIRALpre, which utilizes EVO as its GFM, performs much better than another backbone, and gains nearly 2% improvement in F1-score and 4% in accuracy.

The experiment proves that the proposed training framework fits the foundation model and is a catalyst for classification. With the development of the foundation model, other better-trained models with advanced architecture will appear and VIRALpre can serve to facilitate sequence classification in the future.

## 4 Discussion

With the development of genomic research and foundation model, more and more genome sequences will be discovered, and more powerful models will be proposed. For virus identification, a significant and meaningful task, specificity and precision of tools are highly considered. Nowadays, based on the newest and strongest genomic foundation model, EVO, VIRALpre is proposed and outperforms other tools. Moreover, this K-mer and CNN-based plug-and-play module is raised to manage sequence classification tasks, which acts as a helper to improve the competence of various foundation models and improve the accuracy of the result. Since its simplicity and effectiveness, VIRALpre is promising in the future to facilitate this task with the newly raised foundation model.

Besides binary classification for viruses and other species, many tasks are waiting to be solved using our proposed pipeline. For instance, the classification of single-stranded DNA(ssDNA) phages or proviruses and other viral contigs, a fine-grained task in virus classification, is needed to proceed. What is more, the classification of plasmids, which influences horizontal gene [9] transfer in ecosystems, is urgent to conduct.

There are other points that may help to contribute a more robust tool. For data, only RefSeq datasets are used to train our model, which can be extended to improve the ability of our proposed model. For tokenization, the lengths of input encoded sequences are the same and our model is trained to improve the weaknesses of other tools in short contig classification, however, for longer sequences, this process leads to insufficient utilization of EVO ability of long sequence modeling.

